# Identification of a Putative Sensor Protein Involved in Regulation of Vesicle Production by a Hypervesiculating Bacterium, *Shewanella vesiculosa* HM13

**DOI:** 10.1101/2020.11.12.373589

**Authors:** Fumiaki Yokoyama, Tomoya Imai, Wataru Aoki, Mitsuyoshi Ueda, Jun Kawamoto, Tatsuo Kurihara

## Abstract

Bacteria secrete and utilize nanoparticles, called extracellular membrane vesicles (EMVs), for survival in their growing environments. Therefore, the amount and components of EMVs should be tuned in response to the environment. However, how bacteria regulate vesiculation in response to the extracellular environment remains largely unknown. In this study, we identified a putative sensor protein, HM1275, involved in the induction of vesicle production in a hypervesiculating Gram-negative bacterium, *Shewanella vesiculosa* HM13. This protein was predicted to possess typical sensing and signaling domains of sensor proteins, such as methyl-accepting chemotaxis proteins. Comparison of vesicle production between the *hm1275*-disrupted mutant and the parent strain revealed that HM1275 is involved in lysine-induced hypervesiculation. Moreover, HM1275 has sequence similarity to a biofilm dispersion protein, BdlA, of *Pseudomonas aeruginosa* PAO1, and *hm1275* disruption increased the amount of biofilm. Thus, this study showed that the induction of vesicle production and suppression of biofilm formation in response to lysine concentration are under the control of the same putative sensor protein.

## 1 Introduction

Bacterial cells respond to extracellular environments, create multicellular communities, and communicate with others for survival in their growing environments. To communicate, they secrete and utilize nanoparticles, called extracellular membrane vesicles (EMVs) (Schwechheimer and Kuehn, 2015). Therefore, the amount and components of EMVs should be tuned in response to the environment (Orench-Rivera and Kuehn, 2016). EMVs are composed of various components, such as lipids, proteins, nucleic acids, lipopolysaccharides, and peptidoglycans (Bitto and Kaparakis-Liaskos, 2017). Major components of EMVs of Gramnegative bacteria are derived from the outer membrane and periplasm of the cells, while components of the inner membrane and cytoplasm are also found in EMVs from various bacterial species (Toyofuku et al., 2015; Jain and Pillai, 2017; Tan et al., 2018). Owing to the diversity of these components, EMVs have various physiological roles, being involved in biofilm formation, nutrient uptake, defense (acting as decoys against bacteriophages), and intercellular communication related to horizontal gene transfer and pathogenesis (Toyofuku et al., 2015). Moreover, EMVs have been applied to the development of vaccines and drug delivery systems (Jain and Pillai, 2017; Tan et al., 2018). Elucidation of the vesiculation mechanisms is required for the understanding of their physiology and the development of their applications. Although multiple mechanisms of vesicle production have been suggested to occur in various bacteria (Schwechheimer and Kuehn, 2015; Toyofuku et al., 2015), it remains largely unclear how bacteria regulate the production levels of EMVs in response to extracellular environments.

We previously isolated a bacterial strain suitable for studies of EMVs. This strain, *Shewanella vesiculosa* HM13, is a Gram-negative and cold-adapted bacterium that was isolated from horse mackerel intestines (Chen et al., 2020). Vesiculation by this bacterium has two unique features: higher vesicle production than other strains such as *Escherichia coli* and high purity and productivity of a single specific protein in EMVs; these characteristics make *S. vesiculosa* HM13 useful for elucidating the mechanisms of bacterial vesiculation and protein transport to EMVs (Chen et al., 2020; Guida et al., 2020; Kamasaka et al., 2020; Kawano et al., 2020).

In this study, we identified a putative sensor protein in *S. vesiculosa* HM13 EMVs that harbors sensory Per-Arnt-Sim (PAS) domains and a signaling domain of a methyl-accepting chemotaxis protein (MCP). This protein showed sequence similarity to a biofilm dispersion protein, BdlA, of *Pseudomonas aeruginosa* PAO1 (Morgan et al., 2006; Barraud et al., 2009; Petrova and Sauer, 2012). Gene disruption and quantification of EMVs and biofilm showed that this protein is involved in the sensing of lysine (Lys) in the extracellular environment to induce vesicle production and regulate biofilm formation.

## 2 Materials and Methods

### 2.1 Bacterial Strains and Culture Conditions

The strains used in this study are listed in Table 1. A cold-adapted bacterium, *S. vesiculosa* HM13, was isolated from the intestines of horse mackerel, and a rifampicin-resistant mutant of this strain (HM13-Rif^r^) was used as a parental strain (Chen et al., 2020). A mutant (Δ*gspD2*) lacking a protein required for the transport of a major EMV cargo protein, P49, was used to identify minor proteins included in EMVs (Chen et al., 2020). Another mutant lacking a putative sensor protein encoded by *hm1275* was prepared by gene disruption using a pKNOCK plasmid (Table 1). In brief, a linear fragment of pKNOCK, amplified with the primers pKNOCK-1 and −2 (Table 2), was fused with the internal fragment of *hm1275*, amplified with the primers Dhm1275-1 and −2 (Table 2) using a NEBuilder HiFi DNA Assembly Kit (New England BioLabs, Ipswich, MA, USA) according to the manufacturer’s instructions. *S. vesiculosa* HM13-Rif^r^ was conjugated with *E. coli* S17-1/λ*pir* transformed with the plasmid and then selected on a 1.5% lysogeny broth (LB) agar plate with rifampicin (Rif, 50 μg/mL) and kanamycin (Km, 50 μg/mL) to obtain the mutant Δ*hm1275*. The HM13-Rif^r^ strain and all mutants were subcultured from seed culture using fresh media (1:100 dilution rate) without antibiotics. For the complementation assay of *hm1275*, an empty vector-introduced strain, Δ*hm1275*/p, and the complemented strain, Δ*hm1275*/p*hm1275*, were prepared as described below in the section “Construction of Complemented Strain”. After cultivation, the mutants and empty vector-introduced/complemented strains were plated onto a 1.5% LB agar plate with Rif (50 μg/mL) and Km (50 μg/mL), or with Rif, Km, and chloramphenicol (Cm, 30 μg/mL), respectively, to confirm the maintenance of the plasmid and the plasmid-derived sequence in the genome.

**Table 1.**
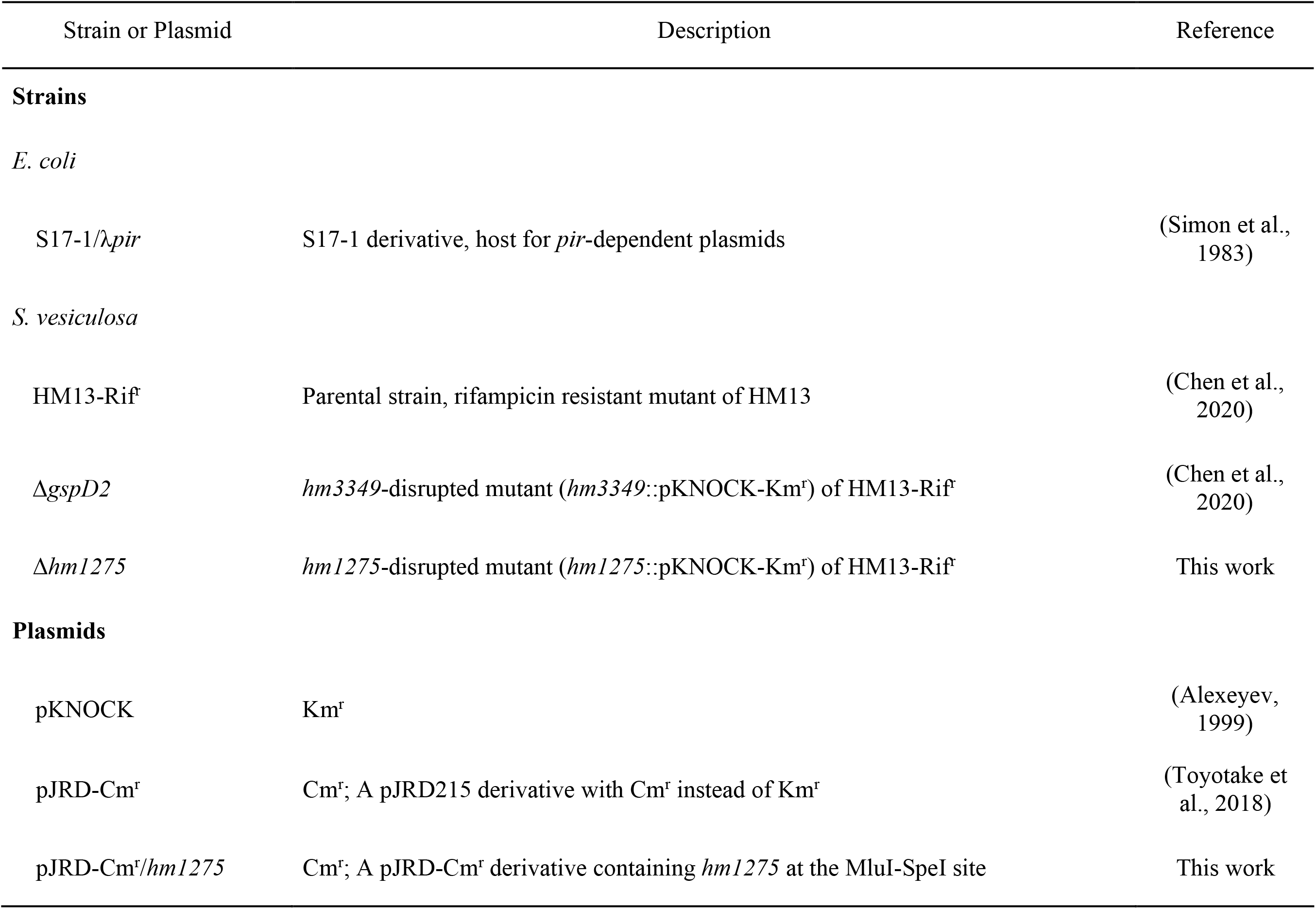
Bacterial strains and plasmids used in this study.

| Strain or Plasmid | Description | Reference |
| --- | --- | --- |
| **Strains** |  |  |
| *E. coli* |  |  |
| S17-1/λ*pir* | S17-1 derivative, host for *pir*-dependent plasmids | (Simon et al., 1983) |
| *S. vesiculosa* |  |  |
| HM13-Rif<sup>r</sup> | Parental strain, rifampicin resistant mutant of HM13 | (Chen et al., 2020) |
| Δ*gspD2* | *hm3349*-disrupted mutant (*hm3349::pKNOCK-Km<sup>r</sup>*) of HM13-Rif<sup>r</sup> | (Chen et al., 2020) |
| Δ*hm1275* | *hm1275*-disrupted mutant (*hm1275::pKNOCK-Km<sup>r</sup>*) of HM13-Rif<sup>r</sup> | This work |
| **Plasmids** |  |  |
| pKNOCK | Km<sup>r</sup> | (Alexeyev, 1999) |
| pJRD-Cm<sup>r</sup> | Cm<sup>r</sup>; A pJRD215 derivative with Cm<sup>r</sup> instead of Km<sup>r</sup> | (Toyotake et al., 2018) |
| pJRD-Cm<sup>r</sup>/*hm1275* | Cm<sup>r</sup>; A pJRD-Cm<sup>r</sup> derivative containing *hm1275* at the MluI-SpeI site | This work |

**Table 2.**
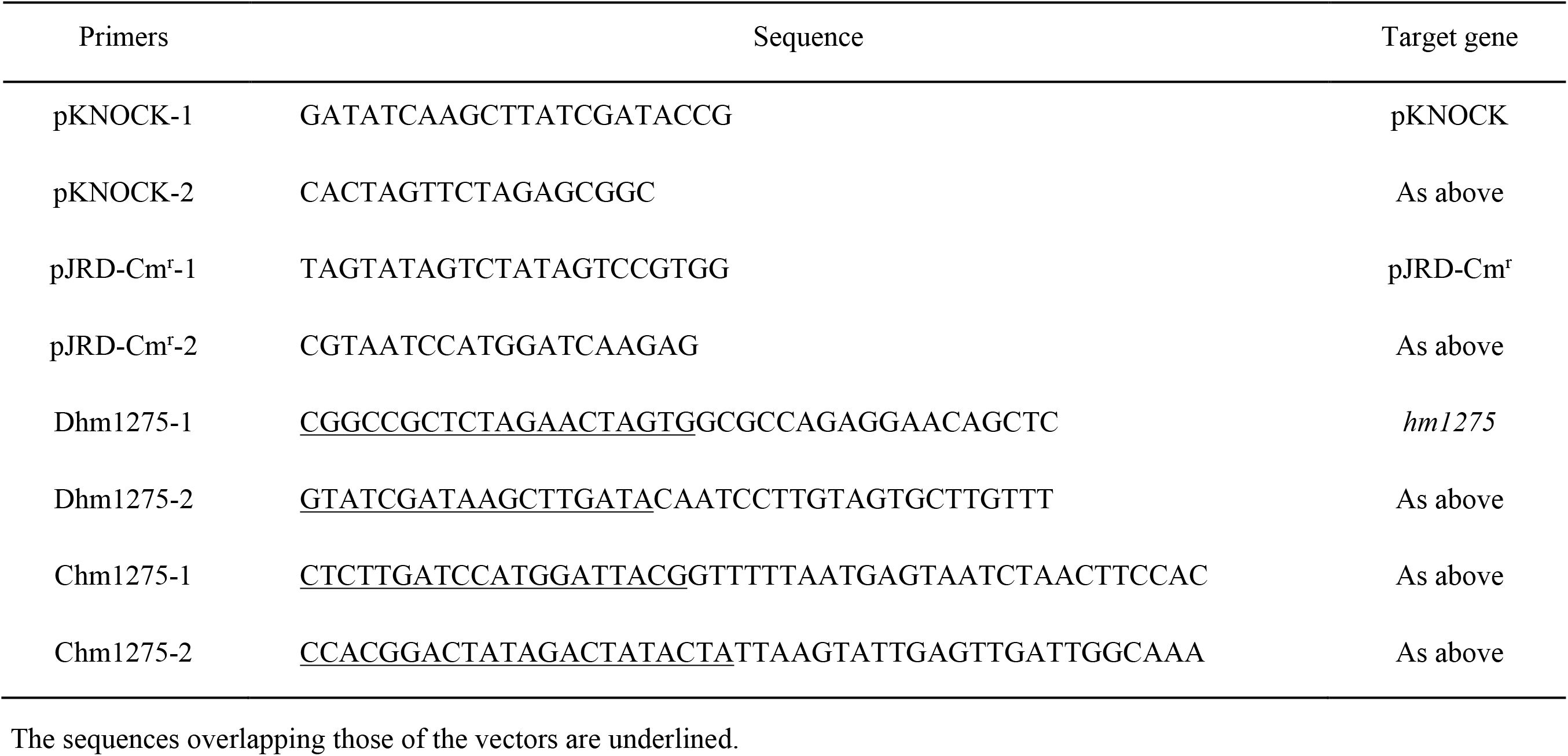
Primers used in this study.

To investigate the effects of the growth environment on vesiculation, three media were used: LB, modified DSMZ medium 79 (M79 medium) (Papa et al., 2006), and Bacto Marine Broth (MB) (Difco, Detroit, MI, USA). The M79 medium contained 1 g/L KH_2_PO_4_, 1 g/L NH_4_NO_3_, 10 g/L NaCl, 0.2 g/L MgSO_4_ · 7H_2_O, 10 mg/L FeSO_4_, 10 mg/L CaCl_2_ · 2H_2_O, and 0.5% w/v casamino acid (CA). M79 medium supplemented with additional CA or Lys (hereafter, M79 + CA or M79 + Lys, respectively) was also used to investigate the effects of amino acids on vesiculation. HM13-Rif^r^ and Δ*hm1275* were grown at 18°C and 180 rpm in a Bio Shaker BR-43FL (Taitec, Saitama, Japan) until the cultures reached the early stationary phase. The optical density at 600 nm (OD_600_) was measured with a UV-visible spectrophotometer (UV-2450, Shimadzu, Kyoto, Japan). The growth curve was obtained using a rocking incubator TVS062CA (ADVANTEC, Tokyo, Japan) at 18°C and 90 rpm.

### 2.2 Construction of Complemented Strain

A DNA fragment including the estimated promoter region and coding region of *hm1275* was obtained byPCR with Q5 High-Fidelity DNA Polymerase (New England BioLabs) using *S. vesiculosa* HM13 genomic DNA as a template and the primers Chm1275-1 and −2 (Table 2). The resulting DNA fragment and linearized pJRD-Cm^r^ (Toyotake et al., 2018), prepared byPCR with the primers pJRD-Cm^r^-1 and −2 (Table 2), were fused to construct pJRD-Cm^r^/*hm1275* using a NEBuilder HiFi DNA Assembly Kit, according to the manufacturer’s instructions. Δ*hm1275* was conjugated with *E. coli* S17-1/λ*pir*, transformed with pJRD-Cm^r^/*hm1275* or the empty vector pJRD-Cm^r^, then selected on a 1.5% LB agar plate with Rif (50 μg/mL), Km (50 μg/mL), and Cm (30 μg/mL) to obtain a complemented strain and the empty vector-introduced strain, Δ*hm1275*/p*hm1275* and Δ*hm1275*/p, respectively.

### 2.3 Isolation of EMVs by Ultracentrifugation

EMVs from *S. vesiculosa* HM13 were collected from cultures at the early stationary phase, unless otherwise stated, according to a previously described method (Yokoyama et al., 2017) with slight modifications. In brief, the cells were pelleted by centrifugation at 6,800 × *g* and 4°C for 10 min. The supernatant was centrifuged at 13,000 × *g* and 4°C for 15 min to remove the remaining bacterial cells. The supernatant was filtered through a 0.45-μm pore polyethersulfone (PES) filter to remove the remaining debris. EMVs were obtained by ultracentrifugation from 4 mL of filtrate at 100,000 × *g* (average centrifugal force) and 4°C for 2 h with an OptimaX centrifuge (Beckman Coulter, Brea, CA, USA). The pellets were resuspended in 400 μL of Dulbecco’s phosphate-buffered saline (DPBS) (Manning and Kuehn, 2011) and used as EMVs in the following experiments. The post-ultracentrifuged supernatant without EMVs was kept as a post-vesicle fraction (PVF) for subsequent experiments.

### 2.4 Vesicle Quantification by Lipid Staining

Lipids in the EMV fractions were quantified with a lipophilic fluorescent molecule, FM4-64 (Molecular Probes/ThermoFisher, Waltham, MA, USA). The fluorescence intensities were divided by the OD_600_ values of the cultures for normalization to quantify vesicle production, according to previously described methods (Yokoyama et al., 2017). To examine the effects of Lys concentration on vesicle production, EMVs of HM13-Rif^r^, statically cultured at 18°C for three days in 5 mL of M79 media with different Lys concentrations, were quantified as described above. The maximum concentration of Lys in this study (2.6 g/L) is supposed to be physiologically relevant and comparable to the one found in the intestine of horse mackerel, the host of *S. vesiculosa* HM13, which consumes krill as its main diet (Huntley et al., 1994; Chen et al., 2009; Kolmakova and Kolmakov, 2019).

### 2.5 Protein Identification by Peptide Mass Fingerprinting

Proteins associated with EMVs (600 μg) of Δ*gspD2* aerobically cultured in LB were identified by two-dimensional polyacrylamide gel electrophoresis (2D-PAGE) and peptide mass fingerprinting, according to previously described methods (Park et al., 2012) with slight modifications. The peptide spectra of each protein spot were subjected to MASCOT search (Matrix Science, London, UK) against the *S. vesiculosa* HM13 protein database. The identified proteins were profiled using BLAST^1^ (Altschul et al., 1997) and HHpred^2^ (Soding et al., 2005). The localization of the identified proteins were predicted using PSORTb version 3.0.2^3^ (Yu et al., 2010) and are listed in Table 3.

**Table 3.**
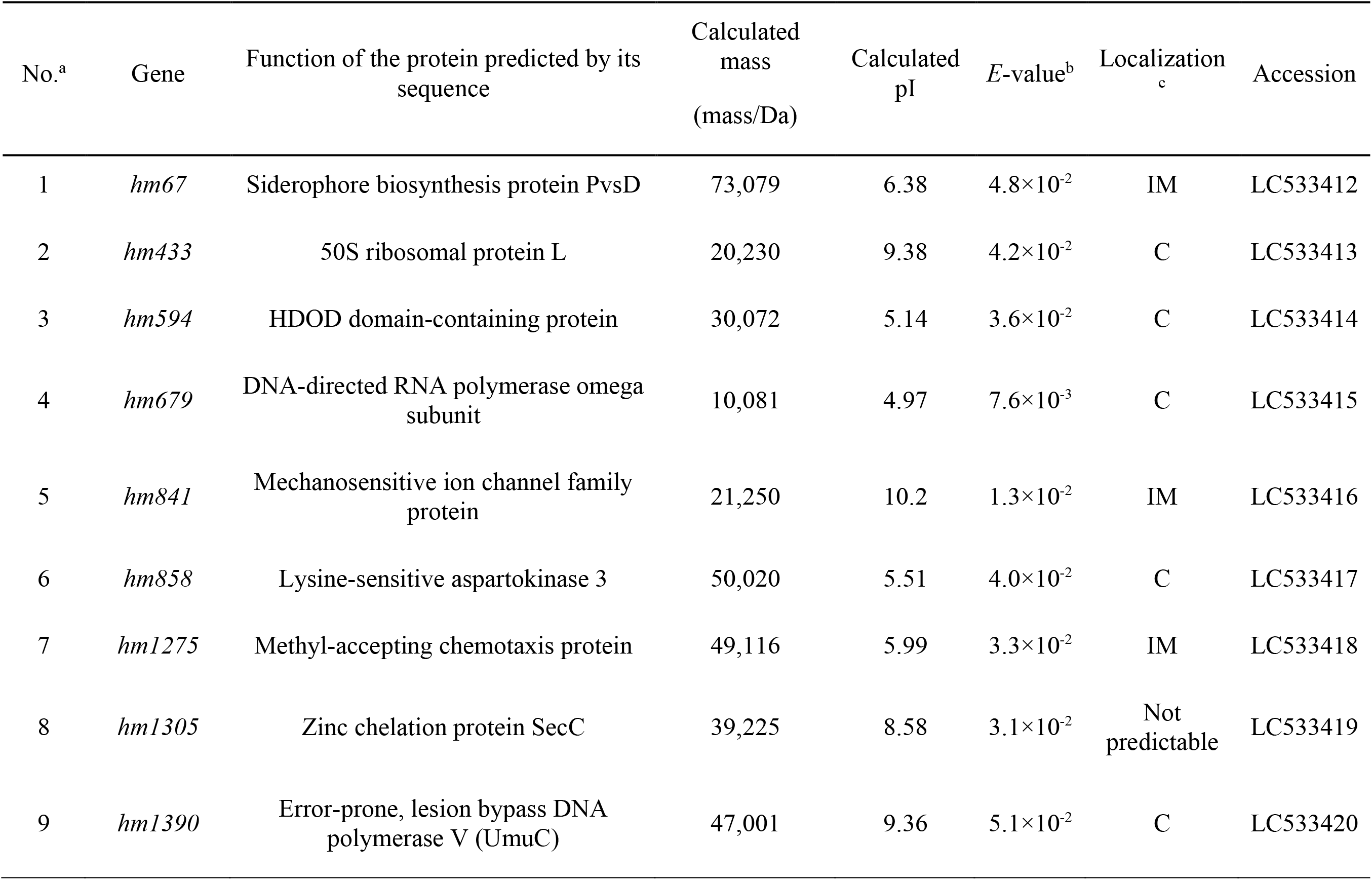
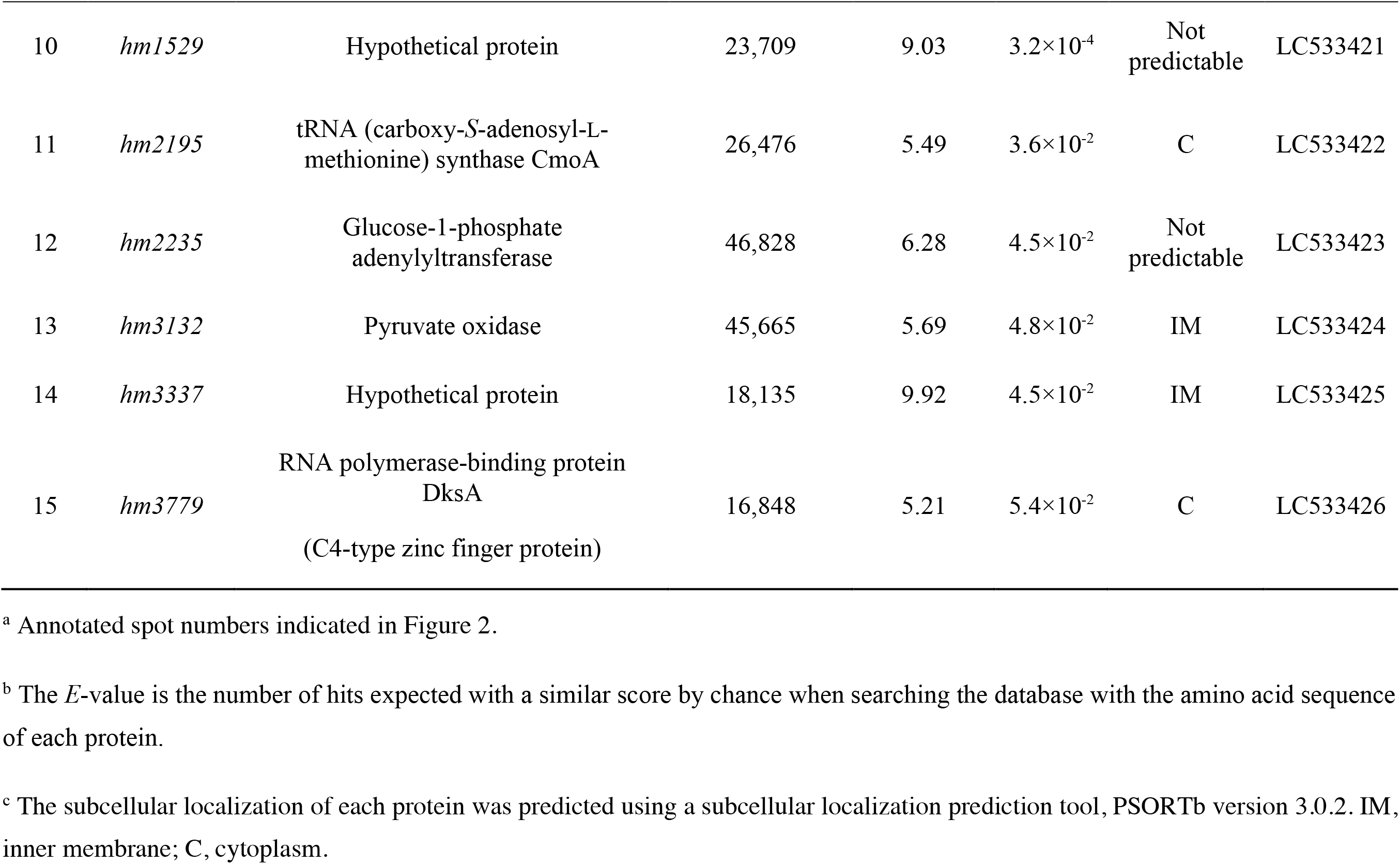
The identified proteins in EMVs from Δ*gspD2*.

| No.<sup>a</sup> | Gene | Function of the protein predicted by its sequence | Calculated mass<br>(mass/Da) | Calculated pI | E-value<sup>b</sup> | Localization<sup>c</sup> | Accession |
| --- | --- | --- | --- | --- | --- | --- | --- |
| 1 | *hm67* | Siderophore biosynthesis protein PvsD | 73,079 | 6.38 | 4.8×10<sup>-2</sup> | IM | LC533412 |
| 2 | *hm433* | 50S ribosomal protein L | 20,230 | 9.38 | 4.2×10<sup>-2</sup> | C | LC533413 |
| 3 | *hm594* | HDOD domain-containing protein | 30,072 | 5.14 | 3.6×10<sup>-2</sup> | C | LC533414 |
| 4 | *hm679* | DNA-directed RNA polymerase omega subunit | 10,081 | 4.97 | 7.6×10<sup>-3</sup> | C | LC533415 |
| 5 | *hm841* | Mechanosensitive ion channel family protein | 21,250 | 10.2 | 1.3×10<sup>-2</sup> | IM | LC533416 |
| 6 | *hm858* | Lysine-sensitive aspartokinase 3 | 50,020 | 5.51 | 4.0×10<sup>-2</sup> | C | LC533417 |
| 7 | *hm1275* | Methyl-accepting chemotaxis protein | 49,116 | 5.99 | 3.3×10<sup>-2</sup> | IM | LC533418 |
| 8 | *hm1305* | Zinc chelation protein SecC | 39,225 | 8.58 | 3.1×10<sup>-2</sup> | Not predictable | LC533419 |
| 9 | *hm1390* | Error-prone, lesion bypass DNA polymerase V (UmuC) | 47,001 | 9.36 | 5.1×10<sup>-2</sup> | C | LC533420 |
| 10 | *hm1529* | Hypothetical protein | 23,709 | 9.03 | $3.2 \times 10^{-4}$ | Not<br>predictable | LC533421 |
| 11 | *hm2195* | tRNA (carboxy-S-adenosyl-L-methionine) synthase CmoA | 26,476 | 5.49 | $3.6 \times 10^{-2}$ | C | LC533422 |
| 12 | *hm2235* | Glucose-1-phosphate<br>adenyltransferase | 46,828 | 6.28 | $4.5 \times 10^{-2}$ | Not<br>predictable | LC533423 |
| 13 | *hm3132* | Pyruvate oxidase | 45,665 | 5.69 | $4.8 \times 10^{-2}$ | IM | LC533424 |
| 14 | *hm3337* | Hypothetical protein | 18,135 | 9.92 | $4.5 \times 10^{-2}$ | IM | LC533425 |
| 15 | *hm3779* | RNA polymerase-binding protein<br>DksA<br>(C4-type zinc finger protein) | 16,848 | 5.21 | $5.4 \times 10^{-2}$ | C | LC533426 |

<sup>a</sup> Annotated spot numbers indicated in Figure 2.
<sup>b</sup> The *E*-value is the number of hits expected with a similar score by chance when searching the database with the amino acid sequence of each protein.
<sup>c</sup> The subcellular localization of each protein was predicted using a subcellular localization prediction tool, PSORTb version 3.0.2. IM, inner membrane; C, cytoplasm.

### 2.6 Quantification of Biofilm by Crystal Violet Staining

The cells attached to the solid surface were stained with crystal violet (CV) to quantify the amount of biofilm, according to previously described methods (Thormann et al., 2004) with slight modifications. In brief, the cells were grown statically in 150 μL of M79 medium with CA or amino acid supplementation in a 96-well U-bottom plate (Delta Lab, Barcelona, Spain) at the optimal growth temperature for HM13-Rif^r^ (18°C) for three days. After cultivation, the floating cells were removed, and the remaining cells attached to the plate were washed twice with 170 μL of water. After drying the plate at room temperature for approximately 5 min, 170 μL of 0.1% CV aq. was added to the wells and incubated at room temperature for 30 min. After the staining solution was removed, the wells were washed twice with 300 μL of water. Then, the cells were destained with 170 μL of 95% ethanol (Nacalai Tesque, Kyoto, Japan) by incubation at room temperature for 30 min. One hundred microliters of the destaining solution were applied to a 96-well flat-bottom plate (Delta Lab) to measure the absorbance at 570 nm with a plate reader (SpectraMax 190, Molecular Devices, San Jose, CA, USA).

### 2.7 Biofilm Observation by Scanning Electron Microscopy

To observe the fine structure of the biofilm formed at the air-liquid interface, the biofilm formed on a glass strip was subjected to scanning electron microscopy (SEM), according to previously described methods (Iwamoto et al., 2007) with slight modifications. In brief, cells in 500 μL of M79 medium with 0.5% CA containing a glass strip along the wall of a 24-well plate (AGC, Tokyo, Japan) were fixed with 2% glutaraldehyde (Wako Pure Chemical Industries, Osaka, Japan), stained with OsO_4_ (Nisshin EM, Tokyo, Japan), subjected to solvent exchange, and lyophilized overnight. The cells on the glass strip were then coated with platinum (approximately 2 nm) using an auto-fine coater (JEC-3000, JEOL, Tokyo, Japan), and observed with a field-emission SEM, JSM-7800F Prime (JEOL), at an acceleration voltage of 5 kV, according to previously described methods (Iwamoto et al., 2007).

### 2.8 Biofilm Observation by Confocal Laser Scanning Microscopy

Biofilms of HM13-Rif^r^ and the mutant were observed by confocal laser scanning microscopy (CLSM), according to previously described methods (Sabaeifard et al., 2017) with slight modifications. In brief, biofilms on a glass-base dish (AGC), obtained by static cultivation at 18°C for three days in 2 mL of M79 medium supplemented with 0.5% CA, were washed with DPBS twice, then stained with 1000-fold diluted propidium iodide and Syto9 (LIVE/DEAD^®^ BacLight^™^ Bacterial Viability Kit, Molecular Probes/ThermoFisher) at room temperature in darkness for 15 min. The biofilms were observed at the air-liquid interface by CLSM using a 60× objective (FV3000, Olympus, Tokyo, Japan) with an excitation laser at 488 nm.

### 2.9 Quantification and Visualization of Biofilm with BiofilmQ

The biofilm images from three frames obtained by CLSM were quantified, and representative images were modeled with the biofilm analysis software BiofilmQ^4^, and the image software ParaView^5^, according to previously described methods (Hartmann et al., 2019).

### 2.10 Protein Identification by Proteome Analysis

Cells of *S. vesiculosa* HM13, statically cultured in 5 mL of M79 media with 0.26 and 2.6 g/L Lys at 18°C for three days, were homogenized by sonication in lysis buffer (7 M urea, 2 M thiourea, 2% 3-[(3-cholamidopropyl)-dimethylammonio]-1-propanesulfonate, 10 mM dithiothreitol, and 50 mM Tris-HCl (pH 7)). The homogenate was centrifuged at 20,000 × *g* at 4°C for 20 min to remove any remaining debris. The solvent was exchanged with 100 mM triethylammonium bicarbonate (pH 8.5) with Amicon Ultra-0.5 (10-MWCO, Millipore, Billerica, MA, USA). Tris(2-carboxyethyl)phosphine (0.2 M) was added to the sample, which was then incubated at 55°C for 60 min. For alkylation, 375 mM 2-iodoacetamide was added, and the sample was incubated at room temperature in darkness for 30 min. Then, cold acetone was added to the sample, which was then incubated at −30°C for 3 h. Next, the sample was centrifuged at 4°C and 20,300 × *g* for 15 min, dried at room temperature, and then suspended in 50 mM triethylammonium bicarbonate. The proteins in the sample were digested with 10 μg/mL sequence grade modified trypsin (Promega, Madison, WI, USA) at 37°C overnight. The digested proteins were labeled by incubation with TMT Mass Tagging Kits and Reagents (Thermo Fischer Scientific, Rockford, IL, USA) in dehydrated acetonitrile at room temperature for 1 h. Then, 5% hydroxylamine was added to the sample, which was then incubated at room temperature for 15 min. Finally, the sample was frozen with liquid nitrogen and stored at −80°C for proteome analysis.

The proteins in the sample were subjected to tandem mass tag-labeling proteomic analysis, according to previously described methods (Tsuji et al., 2020). Proteins were identified by MASCOT search against the *S. vesiculosa* HM13 protein database. The fold-change values were determined as peptide abundance ratios between the cells grown with 0.26 and 2.6 g/L Lys. The global median normalization method was used to normalize the amount of tryptic digests subjected to the analysis. Changes with *p*-values lower than 0.05 were considered statistically significant, and the corresponding genes were annotated by BLAST search and are listed in Table 4.

**Table 4.**
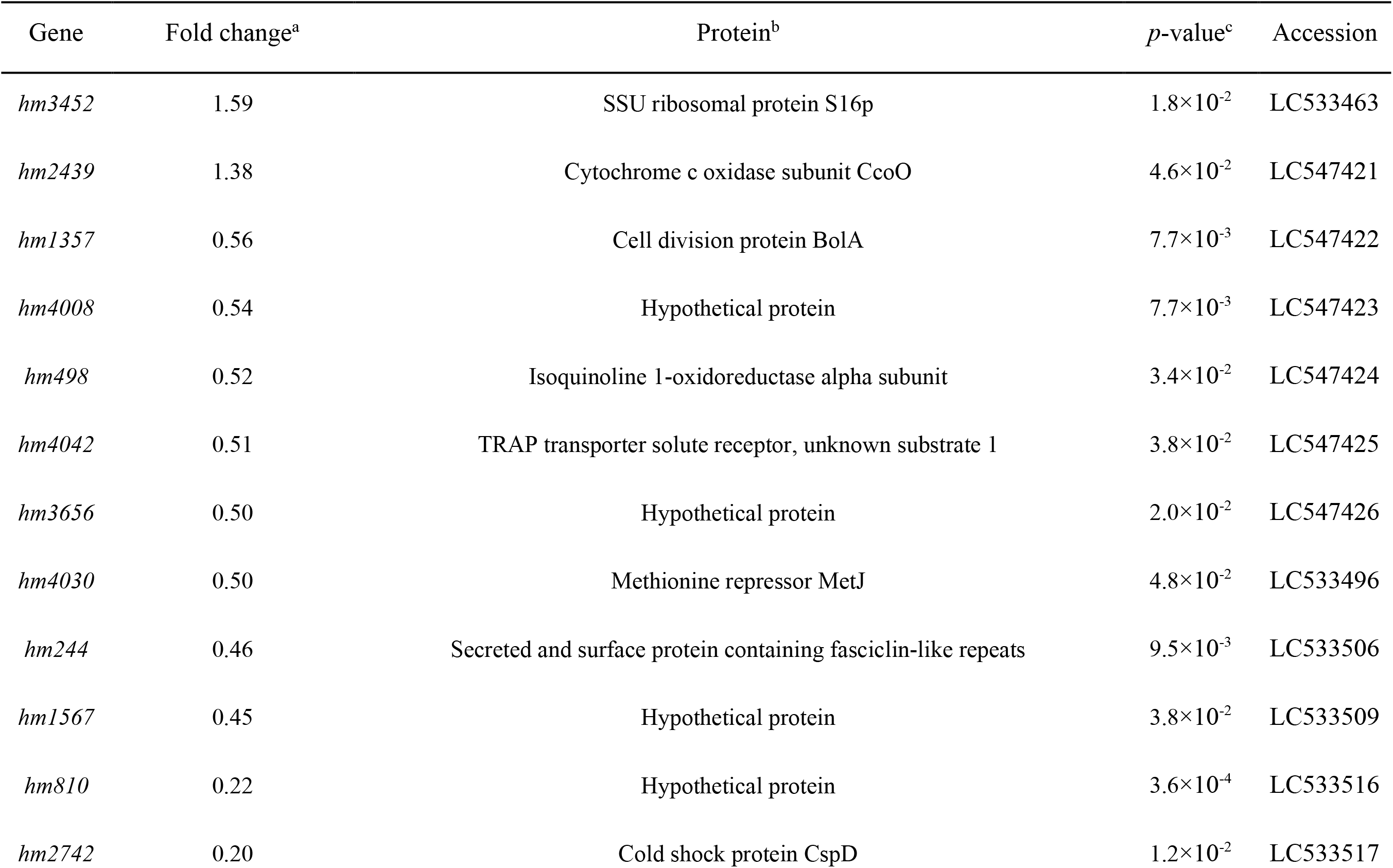
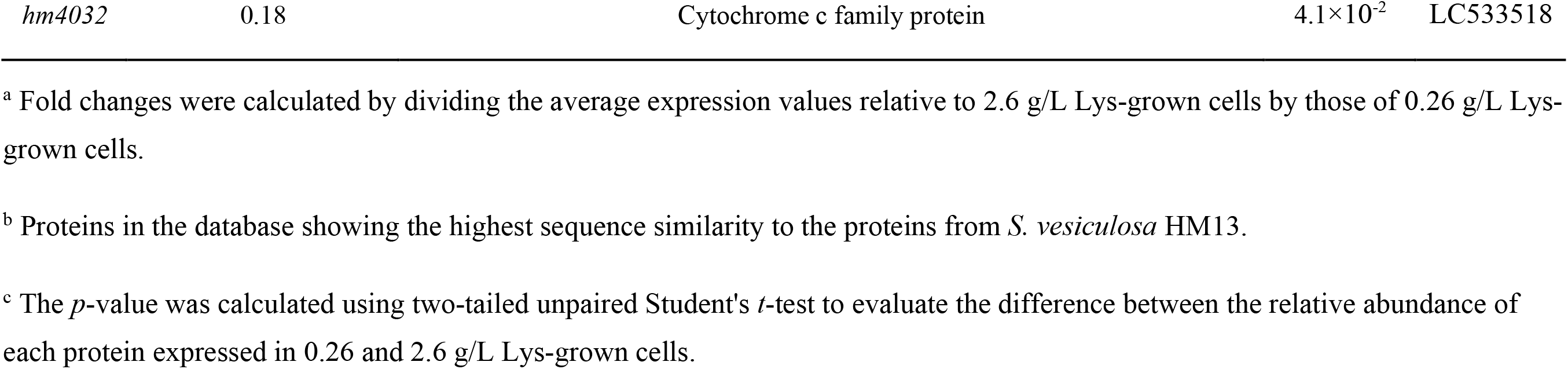
Change of protein expression in response to Lys concentration.

## 3 Results

### 3.1 Variation of Vesiculation Depending on Extracellular Environment

To investigate whether *S. vesiculosa* HM13 regulates vesiculation in response to the extracellular environment, EMVs of this strain cultured in different media were characterized. We used the following three culture media: the rich nutrient LB, a minimal synthetic M79 medium (Papa et al., 2006), and MB, which is generally used for cultivating heterotrophic marine bacteria. The amount of EMVs at the early stationary phase (OD_660_ ≈ 2.5 in LB and 2.0 in M79 medium and MB) (Supplemental Figure S1A) was quantified by lipid staining (Figure 1), which showed that this strain cultured in the nutrient-poor conditions of M79 medium produced fewer EMVs than in the rich nutrient LB, and the amount of EMVs in MB was much lower than that in the other media. Consistent with a previous study (Chen et al., 2020), this strain cultured in LB produced a single major protein, P49, in the EMV fraction, while other proteins were barely visible in both the EMV fraction and the PVF, the latter consisting of the supernatant obtained after ultracentrifugation to remove EMVs (Supplemental Figure S1B). Conversely, in M79 medium and MB, P49 was detected in both EMV fractions and PVFs (Supplemental Figure S1B).

**Figure 1.**
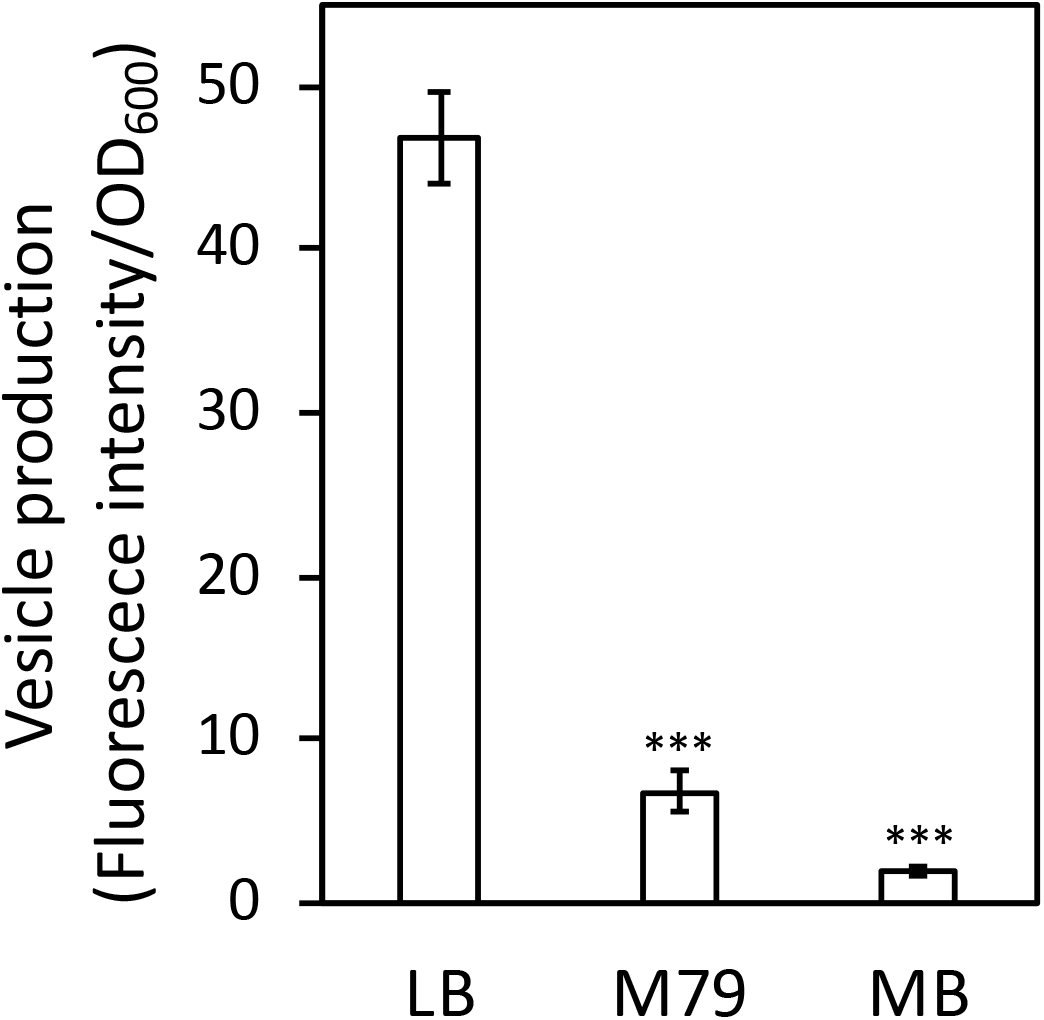
Vesicle productivity of *S. vesiculosa* HM13 in different media. EMVs of *S. vesiculosa* HM13 cultured in different media were quantified by lipid staining with a lipophilic fluorescent molecule, FM4-64. The data are the means ± standard errors of the values from three independent batches. Statistical analysis was performed using two-tailed unpaired Student’s *t*-test. *** indicates*p* < 0.01, compared with LB.

### 3.2 Identification of a Sensor Protein in EMVs

To further characterize EMVs and obtain insights into the mechanism of their biogenesis, proteins in EMVs of the *gspD2*-disrupted mutant (Δ*gspD2*) were identified by gel-based proteomics. For this experiment, Δ*gspD2*, which lacks a putative outer membrane conduit for transport of P49 to EMVs, was used as the disruption of *gspD2* does not impair the production of EMVs (Chen et al., 2020; Kamasaka et al., 2020) and facilitates the identification of minor cargo proteins of EMVs owing to the absence of the major cargo protein P49. Indeed, in EMVs from the parental strain, the amount of P49 is much higher than that of other proteins, and this phenomenon prevents the identification of other proteins by 2D-PAGE owing to the limited protein-loading capacity of the gel.

EMVs from Δ*gspD2* were subjected to 2D-PAGE (Figure 2), and protein spots indicated by the black arrowheads in Figure 2 were identified by peptide mass fingerprinting (Table 3). It is noteworthy that most of them were predicted to be cytosolic and inner membrane proteins; in fact, proteins estimated to localize in the cytoplasm and inner membranes are often found in EMVs (Toyofuku et al., 2015; Jain and Pillai, 2017; Tan et al., 2018). This fact may be at least partially explained by the occurrence of outer-inner membrane vesicles (O-IMVs) secreted by various Gram-negative bacteria (Pérez-Cruz et al., 2013, 2015, 2016).

**Figure 2.**
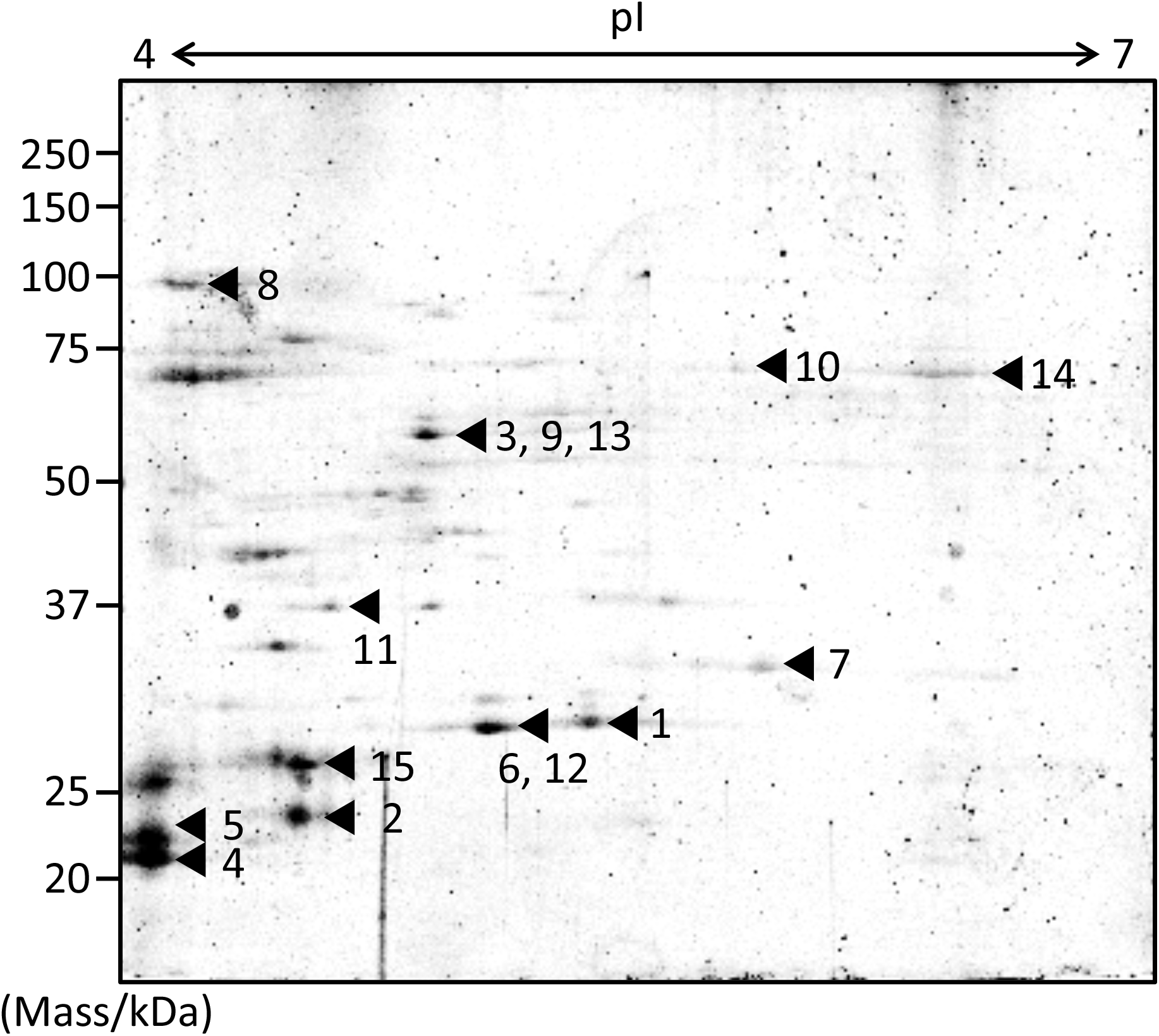
Protein profiling of EMVs. Proteins extracted from the EMVs produced by Δ*gspD2* were subjected to 2D-PAGE and peptide mass fingerprinting. The black arrowheads and the following numbers indicate the identified proteins listed in Table 3. The spot numbered 7 indicates HM1275.

Among these proteins, that encoded by *hm1275* (spot #7 in Figure 2) was profiled using BLAST (Altschul et al., 1997) and HHpred (Soding et al., 2005), which showed that this protein has structural features similar to those of a typical bacterial MCP, which carries a PAS domain to sense signals (Henry and Crosson, 2011) and an MCP signaling domain that interacts with other downstream proteins (Supplemental Table S1) (Falke et al., 1997; Ud-Din and Roujeinikova, 2017). Although proteins that contain PAS and MCP domains generally play a role in signal transduction followed by chemotaxis (Henry and Crosson, 2011), some of these chemosensory components regulate non-chemotaxis-related functions (Wadhams and Armitage, 2004). For example, MCPs have been reported to be involved not only in motility, but also in other bacterial functions, including virulence and biofilm formation (Choi et al., 2015); thus, experimental data are needed to elucidate the function of HM1275. Since HM1275 was predicted to be a membrane protein using PSORTb (Yu et al., 2010), its properties may have caused its abnormal migration in 2D-PAGE, where its apparent molecular mass differed from the calculated mass reported in Table 3; in fact, this phenomenon also occurred for other membrane proteins (Rath et al., 2009; Rath and Deber, 2013).

### 3.3 Induction of Vesicle Production Mediated by a Putative Sensor Protein, HM1275, in the Presence of a High Concentration of Lys

We then examined whether HM1275 senses signal molecules in the extracellular environment and regulates vesicle production. To facilitate the identification of a signal molecule that affects vesicle production, a totally synthetic and nutrient-poor M79 medium was used for this experiment. Within its components, we focused on amino acids, previously reported as signal molecules for MCPs in bacteria (Springer et al., 1977; Hedblom and Adler, 1980; Hanlon and Ordal, 1994). In particular, we focused on the concentration of Lys, which is abundant in zooplankton (Kolmakova and Kolmakov, 2019). The latter is the most typical diet of horse mackerel (Jardas et al., 2004), from whose intestines *S. vesiculosa* HM13 was isolated (Chen et al., 2020). Therefore, we hypothesized that Lys concentration may be sensed by this strain as a marker of environmental change caused by food intake by the host; thus, we cultivated bacterial cells in media containing Lys at physiological concentrations (up to 2.6 g/L) for subsequent experiments (for details, please refer to the Materials and Methods section).

An *hm1275*-disrupted strain (Δ*hm1275*) was prepared as described in the Materials and Methods section. The growth of this strain was similar to that of its parental strain (HM13-Rif^r^) in M79 + CA (Supplemental Figure S2). Next, these two strains were cultured in modified M79 media supplemented with different concentrations of Lys. The relative vesicle production of HM13-Rif^r^, quantified by lipid staining, increased in a Lys concentration-dependent manner (Figure 3). Similar tendencies were observed by nanoparticle tracking analysis (Supplemental Figure S3A) (Filipe et al., 2010). Moreover, EMVs produced under these conditions had similar mean diameters, although smaller EMVs were more abundant in the presence of 1.3 g/L Lys than in other conditions (Supplemental Figure S3B). Notably, HM13-Rif^r^ cultured in M79 medium with 2.6 g/L Lys produced an approximately four-fold larger number of EMVs compared to the same strain grown in original M79 medium containing 0.26 g/L Lys. However, Lys-induced vesicle production by Δ*hm1275* was significantly lower than that of HM13-Rif. Nevertheless, under all conditions, transmission electron microscopy did not reveal any morphological differences between EMVs from these strains (Supplemental Figure S4).

**Figure 3.**
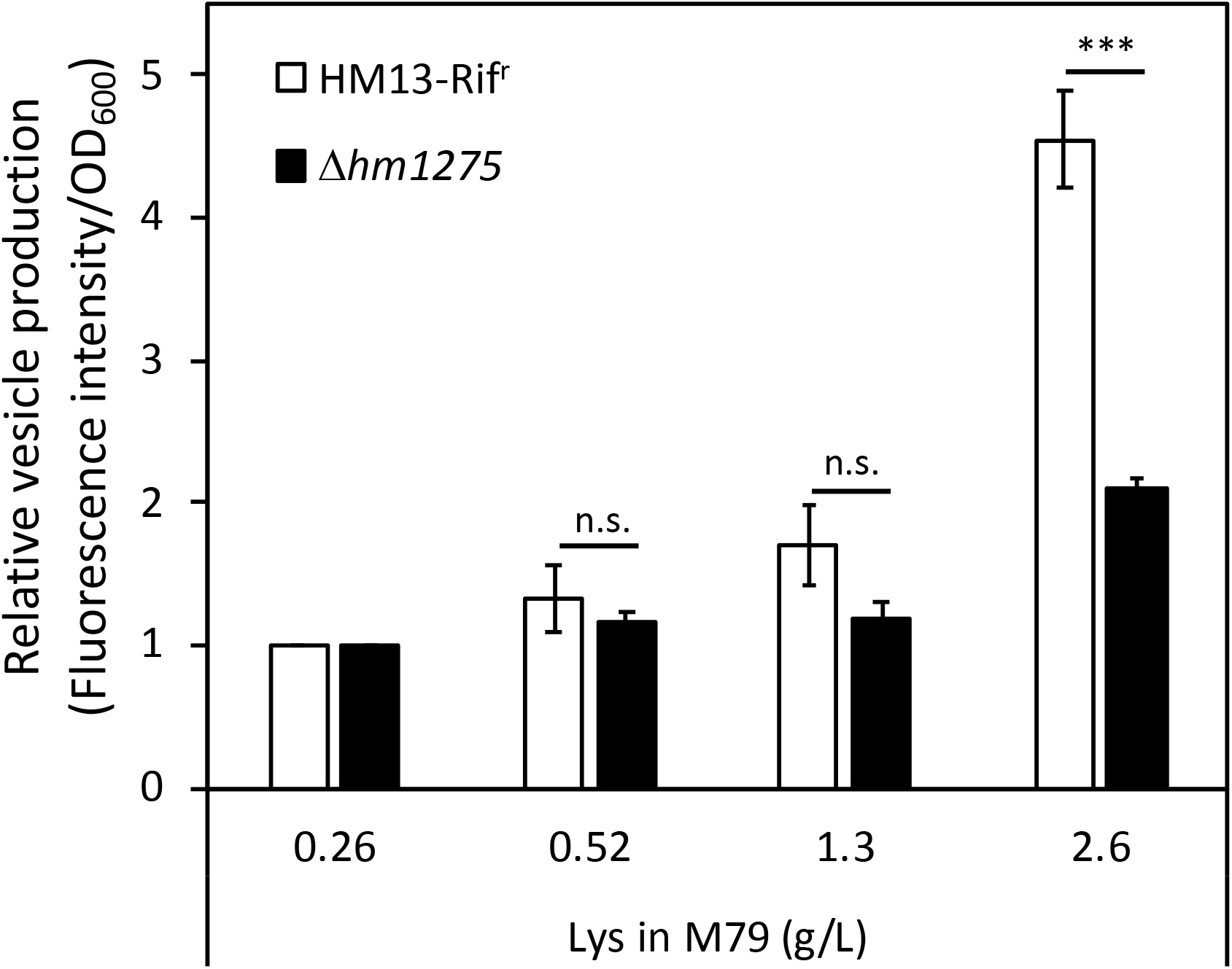
Induced vesicle production of *S. vesiculosa* HM13 by the addition of Lys in a concentration-dependent manner. The vesicle production of HM13-Rif^r^ and Δ*hm1275* cultured at different Lys concentrations was quantified by lipid staining. Each value of vesicle production was divided by the absorbance of cells grown in M79 containing 0.26 g/L Lys to compare relative vesicle production in each condition. The data are the means ± relative standard errors of the values from three independent batches. Statistical analysis was performed using two-tailed unpaired Student’s *t*-test. n.s. and *** indicate *p* ≥ 0.1 and *p* < 0.01, respectively.

To confirm that this phenotype was caused by the disruption of *hm1275*, the complemented strain of Δ*hm1275* (Δ*hm1275*/p*hm1275*) and an empty vector-introduced strain (Δ*hm1275*/p) were used. In M79 media with 2.6 g/L Lys, the relative vesicle production of Δ*hm1275*/p*hm1275* was significantly higher than that of Δ*hm1275*/p (Supplemental Figure S5), suggesting that the disruption of *hm1275* caused the observed phenotype.

### 3.4 Repression of HM1275-Dependent Biofilm Formation

To adapt to the extracellular environment, bacteria accurately control the transition between planktonic and biofilm states (Rossi et al., 2018). Interestingly, pairwise alignment of amino acid sequences with Clustal Omega^6^ (Sievers et al., 2011) revealed that HM1275 has a 38.8% sequence identity to BdlA, a protein related to biofilm dispersion in *P. aeruginosa* PAO1 (accession number: Q9I3S1; Supplemental Figure S6) (Morgan et al., 2006; Barraud et al., 2009; Petrova and Sauer, 2012). This protein plays a role in sensing the extracellular environment and downregulates the accumulation of cyclic-di-GMP to induce biofilm dispersion. This sequence similarity suggests that HM1275 is not only involved in vesicle production, but also in biofilm formation (Bobrov et al., 2005; Thormann et al., 2006). Therefore, biofilm formation by HM13-Rif^r^ and Δ*hm1275* was measured over time by staining with CV (Thormann et al., 2004). We observed that the amount of biofilm in HM13-Rif^r^ decreased in a time-dependent manner, but such a decrease was much less significant in the mutant (Figure 4A).

**Figure 4.**
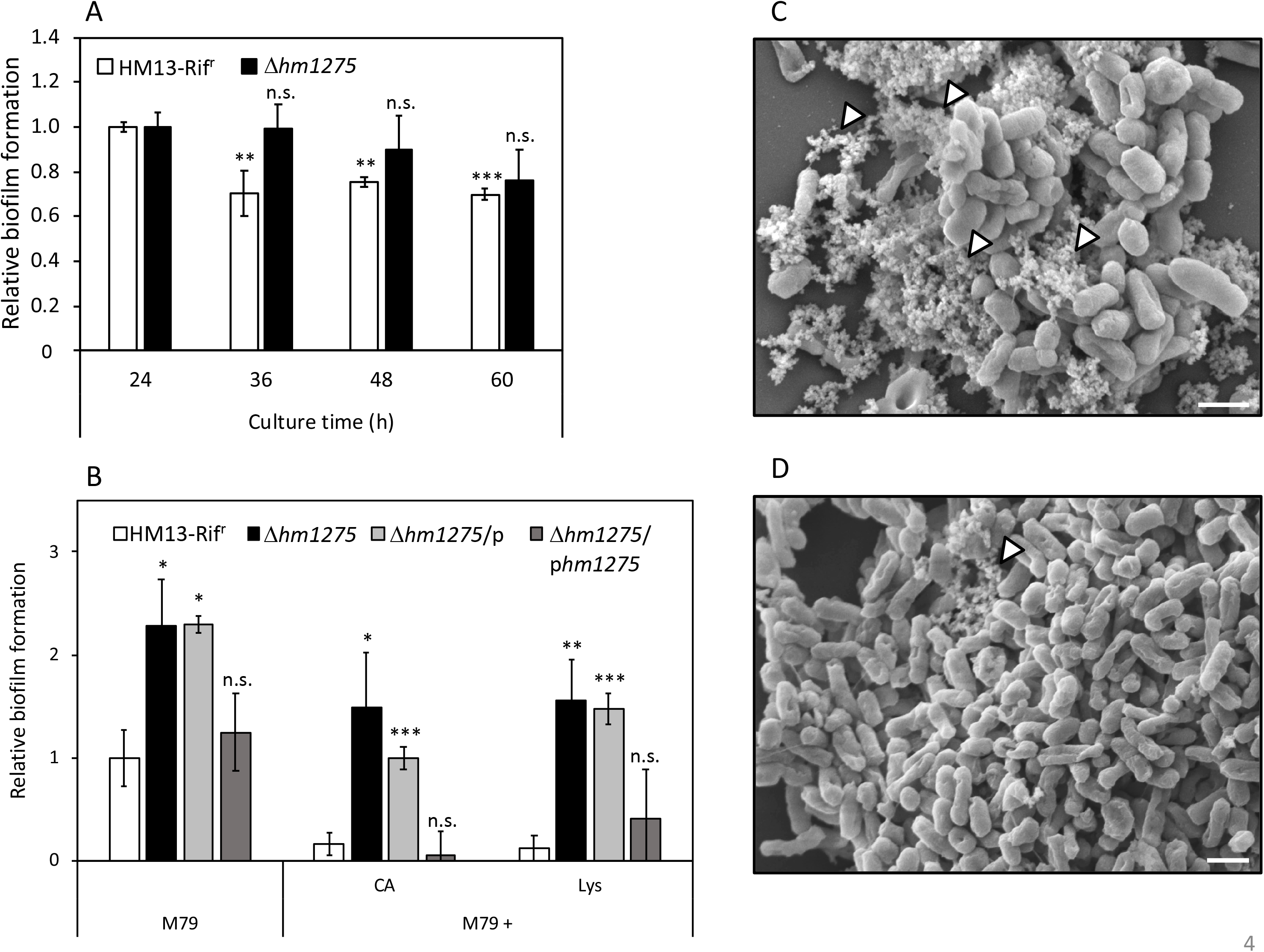
Suppression of biofilm formation of *S. vesiculosa* HM13 by the addition of Lys. (A) The biofilm formation of HM13-Rif^r^ and Δ*hm1275* cultured in M79 + CA was quantified in time-course by staining with CV. (B) Biofilm formation of HM13-Rif^r^ and Δ*hm1275* in modified M79 media was quantified after 72 h of cultivation by staining with CV. The data are the means ± relative standard errors of the values from three independent batches. Statistical analyses for differences between 24 h and other culture times (A) and for differences between HM13-Rif^r^ and other strains in each cultivation condition (B) were performed using two-tailed unpaired Student’s *t*-test. n.s., *, **, and *** indicate *p* ≥ 0.1, and *p* < 0.1, 0.05, and 0.01, respectively. (C and D) The fine structures of biofilm of HM13-Rif^r^ (C) and Δ*hm1275* (D) were observed by SEM. The white arrowheads indicate stackings of EMV-like structures. The bars indicate 1 μm.

To examine whether biofilm formation is controlled by Lys concentration in an HM1275-dependent manner, similar to vesicle production, biofilm formation of HM13-Rif^r^ and Δ*hm1275* cultured in M79 media supplemented with CA and Lys was quantified (Figure 4B). Under all conditions tested, the relative amounts of biofilm from HM13-Rif^r^ were lower than those of Δ*hm1275* (*p* = 0.067, 0.073, and 0.025 for bacteria grown in M79, M79 + CA, and M79 + Lys, respectively) and the empty vector-introduced strain (Δ*hm1275*/p) (*p* = 0.010, 0.0062, and 0.0024 for bacteria grown in M79, M79 + CA, and M79 + Lys, respectively), but were not significantly different from those of the complemented strain (Δ*hm1275*/p*hm1275*). Moreover, the amounts of biofilm from HM13-Rif^r^ decreased to 16.8% upon addition of CA and to 12.0% upon addition of Lys, whereas those of the mutant decreased by a lesser extent upon addition of CA (65.1%) and Lys (68.3%), compared with those in the original M79 medium. These results suggest that HM1275 is involved in Lys-induced suppression of biofilm formation.

Next, we analyzed the fine structures of biofilms of HM13-Rif^r^ and Δ*hm1275* by SEM. Some stackings of EMV-like structures were frequently observed in HM13-Rif^r^ biofilm (white arrowheads in Figure 4C). Conversely, such structures were much less frequently observed in the mutant biofilm (white arrowheads in Figure 4D).

### 3.5 Increased Population of Dead Cells in the Biofilm of Δ*hm1275*

We further investigated the biomass and live/dead cell ratio in biofilms of HM13-Rif^r^ and Δ*hm1275* by CLSM. For HM13-Rif^r^, a small amount of biofilm (red square in Figure 5A) and many single cells attached to the glass bottom of the dish base were observed (Figure 5A) compared with the mutant, for which a large amount of biofilm was observed (Figure 5B). Furthermore, mutant biofilms contained a larger number of dead cells than HM13-Rif^r^ biofilms (Figure 5A and B). In particular, the sectioned images in CLSM and models of biofilm indicated that the HM13-Rif^r^ biofilm included a small number of dead cells in its interior region (Figure 5 A and C), whereas the Δ*hm1275* biofilm included abundant dead cells at the bottom and in its interior region (Figure 5B and D). Biomass quantification of the biofilm of these strains showed that the total biomass of the mutant biofilm was approximately five-fold larger than that of the HM13-Rif^r^ biofilm (Figure 5E). However, the HM13-Rif^r^ biofilm was composed of approximately 80% live and 20% dead cells, while the mutant biofilm was composed of approximately 53% live and 47% dead cells.

**Figure 5.**
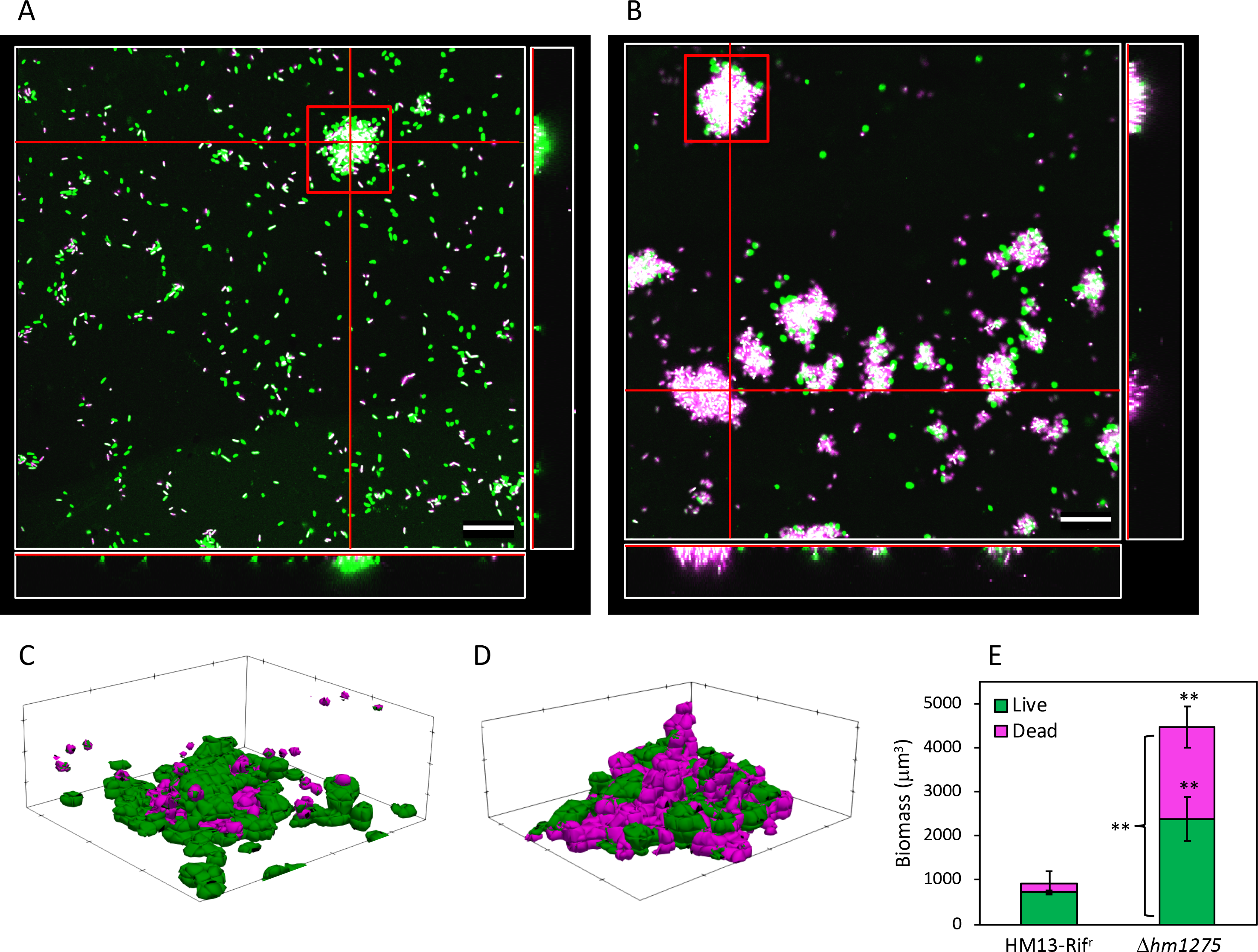
Live/dead cell analysis of surface-associated cells of HM13-Rif^r^ and Δ*hm1275*. Surface-associated cells of HM13-Rif^r^ (A) and Δ*hm1275* (B) were observed by CLSM. Green, magenta, and white colors indicate living and dead cells, and merged colors, respectively. The white boxes at the right and bottom of each image show cross-sections at the vertical and horizontal red lines in the center white box, respectively. The central white box of each image shows a side-cross-section at the red lines in the right and bottom white boxes. The bars indicate 20 μm. (C and D) Models of biofilm of HM13-Rif^r^ (C) and Δ*hm1275* (D) in the red square in A and B were constructed with BiofilmQ and ParaView. (E) The biomass and live/dead ratio in biofilm were quantified with BiofilmQ. The data are the means ± standard errors of the values from three independent batches. Statistical analysis was performed for differences between HM13-Rif^r^ and Δ*hm1275* using two-tailed unpaired Student’s *t*-test. ** indicates *p* < 0.05.12

### 3.6 Comprehensive Identification of Lys-Induced and -Repressed Proteins

To obtain insights into the mechanisms of Lys-induced vesicle production and suppression of biofilm formation mediated by HM1275, proteins up- and downregulated by increasing the concentration of Lys in the medium from 0.26 to 2.6 g/L were identified by shotgun proteomics (Supplemental Figure S7). Proteome analysis covered approximately 35% of all gene products of *S. vesiculosa* HM13. The genes encoding proteins that showed significant differences in expression levels between cells grown with 0.26 g/L and 2.6 g/L Lys were annotated by a BLAST search and are listed in Table 4. These proteins (n = 13) accounted for approximately 1% of the total proteome (n = 1,488). Among these proteins, two proteins, a ribosome subunit and a cytochrome c oxidase subunit (HM3452 and HM2439, respectively), were slightly upregulated (Table 4). On the other hand, 11 downregulated proteins included proteins related to cell division (HM1357), oxide reduction (HM498 and HM4032), transportation (HM4030 and HM4042), cell adhesion (HM244), and DNA replication (HM2742) (Table 4). The possible involvement of these proteins in vesicle production and biofilm formation will be discussed in the Discussion section.

## 4 Discussion

In this study, we found that the putative sensor protein HM1275 of *S. vesiculosa* HM13 is involved in the regulation of vesicle production and biofilm formation. Vesicle production and suppression of biofilm formation were co-induced through HM1275 in response to high Lys concentrations, suggesting a common regulation mechanism shared between these physiological responses.

To explore the mechanism of HM1275-mediated response to extracellular Lys concentration, we identified proteins whose expression levels were significantly changed in response to Lys concentration. When the cells were grown with high concentrations of Lys, HM1357, a homolog of a cell division-related protein, BolA, was downregulated (0.56-fold) compared with its expression in cells grown with low concentrations of Lys (Table 4). BolA regulates the transcript levels of several proteins involved in peptidoglycan synthesis, and its deletion causes loss of integrity of the outer membrane (Freire et al., 2006). Furthermore, the deletion of a gene coding for a BolA-like protein, IbaG, reduces peptidoglycan levels and alters outer membrane lipids, disturbing membrane integrity (Fleurie et al., 2019). Decreased membrane peptidoglycan crosslinking plays a role in vesicle formation in Gram-negative bacteria (Nagakubo et al., 2020). Therefore, it is plausible that the downregulated expression of HM1357, a BolA-like protein of *S. vesiculosa* HM13, reduces the stability of the peptidoglycan network and weakens the linkage between peptidoglycan layers and the outer membrane, thereby inducing membrane blebbing and subsequent vesicle production.

We found that HM1275 is also involved in Lys-induced suppression of biofilm formation. Therefore, cells are also supposed to regulate the expression of proteins related to biofilm formation in response to Lys. BolA is also involved in biofilm formation by facilitating the production of fimbria-like adhesins and curli, and negatively modulating flagellar biosynthesis and swimming capacity (Dressaire et al., 2015). Even in the presence of an abundant biofilm-forming factor, cyclic-di-GMP, the absence of BolA reduced the amount of biofilm (Moreira et al., 2017). Therefore, we speculate that HM1275 senses extracellular signals and downregulates the expression of the BolA-like protein HM1357 to suppress biofilm formation.

We also found that HM244, a homolog of a secreted and surface protein containing fasciclin-like repeats, was 0.46-fold downregulated under high Lys conditions (Table 4). The fasciclin domain is involved in cell attachment across plants, animals, and bacteria (Seifert, 2018). Therefore, the decreased production of HM244 in response to high Lys concentration may decrease biofilm formation by suppressing cell adhesion.

Biofilm formation may play an important role in the survival of *S. vesiculosa* HM13 in its original habitat. This strain was isolated from fish intestines, where nutrients fluctuate depending on the feeding activity of the fish; thus, this bacterium should cope well with nutrient fluctuations for its survival (Pereira and Berry, 2017). When nutrients are scarce in the host intestine, biofilm formation is thought to be beneficial for bacterial survival, as reported for various strains (Falkinham III, 2009; Cherifi et al., 2017). On the other hand, when the host fish ingests food, the digested food is introduced to the intestine, rendering the environment suitable for bacteria to proliferate. Under these conditions, it is likely that *S. vesiculosa* HM13 cells respond to Lys, which is supposed to be abundant in the diet of the host fish, through the sensor protein HM1275, thereby suppressing biofilm formation. Inside and at the bottom of the biofilm, cells are subjected to low oxygen and nutrient availability (Flemming et al., 2016). Consequently, if the cells cannot properly suppress biofilm formation under nutrient-rich conditions, they cannot benefit from such conditions and are expected to die because of the limited resources available in the biofilm (Figure 5) (Cherifi et al., 2017). Therefore, regulation of biofilm formation in a timely manner, probably by HM1275, contributes to the survival of this strain in a changeable environment.

HM1275 was identified as a cargo of EMVs (Figure 2), although it was predicted to localize to the inner membrane by PSORTb (Yu et al., 2010) (Table 3). These results suggest that HM1275 found in the EMV fraction is included in O-IMVs, which display membranes derived from the cell’s inner membrane and are produced by various Gram-negative bacteria as a minor fraction of EMVs (Pérez-Cruz et al., 2013, 2015, 2016). This interpretation is supported by our previous observation that *S. vesiculosa* HM13 produces EMVs with a double membrane-bounded structure, which are supposed to be O-IMVs (Chen et al., 2020). On the other hand, the physiological significance of the occurrence of HM1275 in EMVs is currently unknown. As a putative sensor protein, HM1275 is thought to play a role in the cellular inner membrane, where it performs signaling functions that facilitate EMV production and suppression of biofilm formation. Thus, HM1275 packaged into EMVs may not have an active role, but rather be eliminated from the cells as a result of protein quality control (McBroom and Kuehn, 2006) or released into the culture supernatant as a result of explosive cell lysis (Turnbull et al., 2016). Nevertheless, another hypothesis may also be that EMVs function as a carrier of HM1275 for the transfer of this protein to other cells to facilitate collective cell behavior such as the regulation of biofilm formation, although, to our knowledge, the intercellular transfer of inner membrane proteins has not been demonstrated so far. These hypotheses should be examined in future studies. Although the physiological significance of HM1275 in EMVs remains elusive, the present study revealed that a putative sensor protein is involved in nutrient-responsive coregulation of EMV production and biofilm formation. A possible functional link between EMV production and biofilm formation will also be examined in future studies.

## 5 Conflict of Interest

The authors declare that the research was conducted in the absence of any commercial or financial relationships that could be construed as a potential conflict of interest.

## 6 Author Contributions

FY, JK, and TK designed the study. FY performed the experiments. TI contributed to electron microscopic analysis. WA and MU contributed to proteome analysis. FY, JK, and TK discussed the results and wrote the manuscript. All authors have read and approved the submitted version.

## 7 Funding

This work was supported by research grants from JSPS KAKENHI (JP15H04474, JP17H04598, JP18H02127, and JP18K19178 to TK, and JP16K14885 to JK), the Institute for Fermentation, Osaka (L-2019-2-012 to TK), and JSPS Research Fellowship for Young Scientists (to FY).

## 8 Acknowledgments

We thank Prof. Hiroyuki Yano (Research Institute for Sustainable Humanosphere, Kyoto University) for SEM observation of bacterial biofilm; Prof. Shiroh Futaki and Dr. Kenichi Kawano (Institute for Chemical Research, Kyoto University) for CLSM analysis of bacterial biofilm; Prof. Kazunari Akiyoshi and Dr. Asako Shimoda (Graduate School of Engineering, Kyoto University) for nanoparticle tracking analysis of bacterial vesicles. Transmission electron microscopy was performed in collaboration with the Analysis and Development System for Advanced Materials (ADAM) at the Research Institute for Sustainable Humanosphere, Kyoto University.

## 10 Data Availability Statement

The genes identified in this study have been submitted to the DNA Data Bank of Japan, and the accession numbers listed in Tables 3 and 4 were assigned.

## Footnotes

1 http://www.ncbi.nlm.nih.gov/blast

2 https://toolkit.tuebingen.mpg.de/tools/hhpred

3 http://www.psort.org/psortb/

4 https://drescherlab.org/data/biofilmQ/docs/index.html

5 https://www.paraview.org/

6 https://www.ebi.ac.uk/Tools/msa/clustalo/

## Notes

### Competing Interest Statement

The authors have declared no competing interest.

