## Supplemental for "Identification of a Putative Sensor Protein Involved in Regulation of Vesicle Production by a Hypervesiculating Bacterium, *Shewanella vesiculosa* HM13"

### *Supplementary Material*

#### 1 Supplementary Materials and Methods

##### 1.1 SDS-PAGE

To analyze the proteins associated with EMVs, EMVs and PVF were subjected to trichloroacetic acid (Wako Pure Chemical Industries, Osaka, Japan) precipitation and SDS-PAGE, according to a previously described method (Yokoyama et al., 2017) with slight modifications in the staining step, where CBB was used instead of SYPRO Ruby.

##### 1.2 Vesicle quantification by Nanoparticle Tracking Analysis

The number of EMVs was quantified by nanoparticle tracking analysis according to a previously described method (Shimoda et al., 2016) with slight modifications. In brief, the EMVs of HM13-Rif$^r$ and the mutants were diluted to about $10^8$–$10^9$ particles/mL and analyzed using NanoSight LM10 (NanoSight, Amesbury, UK) with a blue laser at a shutter speed of 1500, camera gain of 500, and threshold of eight for 60 s.

##### 1.3 Morphological analysis of EMVs by Transmission Electron Microscopy

Negative-stained EMVs of *S. vesiculosa* HM13 cultured in different media were observed by transmission electron microscopy (JEM-1400, JEOL, Tokyo, Japan) with a built-in charge-coupled device camera, according to previously described methods (Yokoyama et al., 2017).

#### 3 Supplementary Figures and Tables

##### 3.1 Supplementary Figures

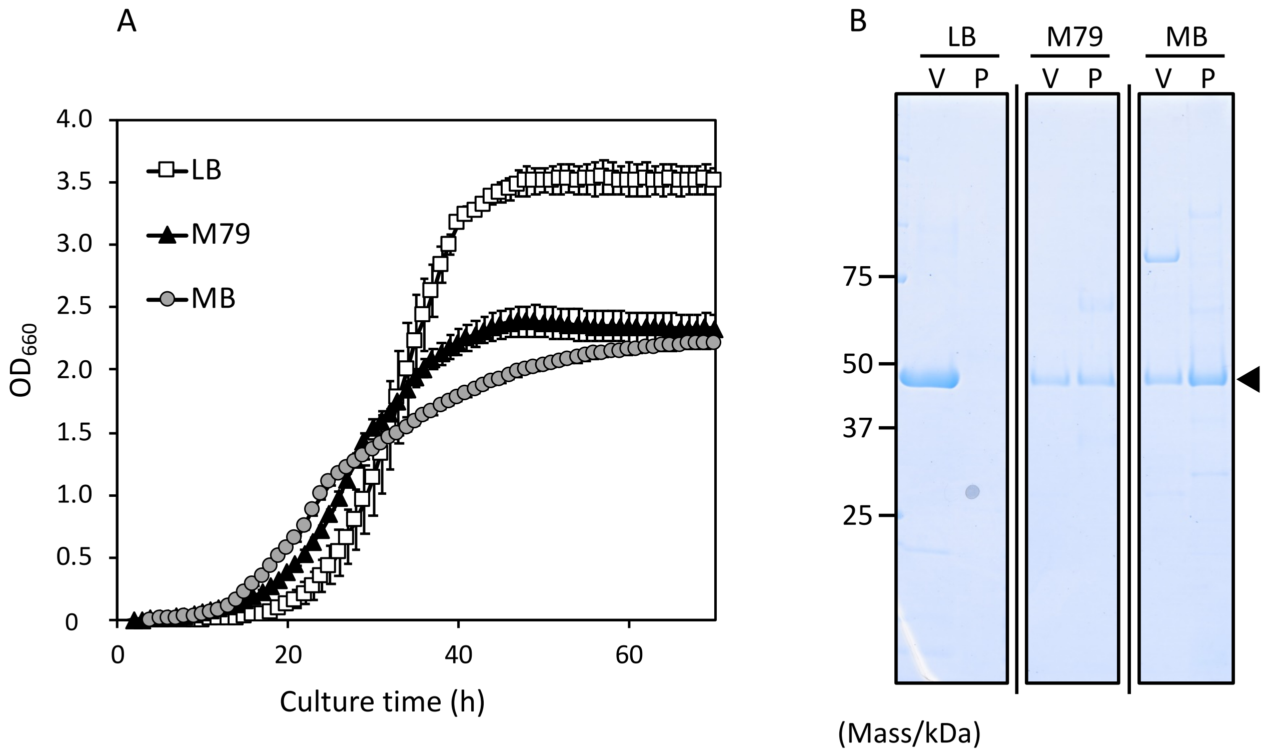


**Figure S1.** Growth patterns and protein compositions of EMVs and PVFs of HM13-Rif$^{r}$ in different media. (A) Growth curves of *S. vesiculosa* HM13. The data are the means ± standard errors of the values from three independent batches. (B) Representative images of two independent batch experiments in which proteins were analyzed by SDS-PAGE. The black arrowhead indicates the position of P49. The leftmost numbers indicate molecular masses of protein standards. “V” and “P” indicate the EMV fraction and PVF, respectively.

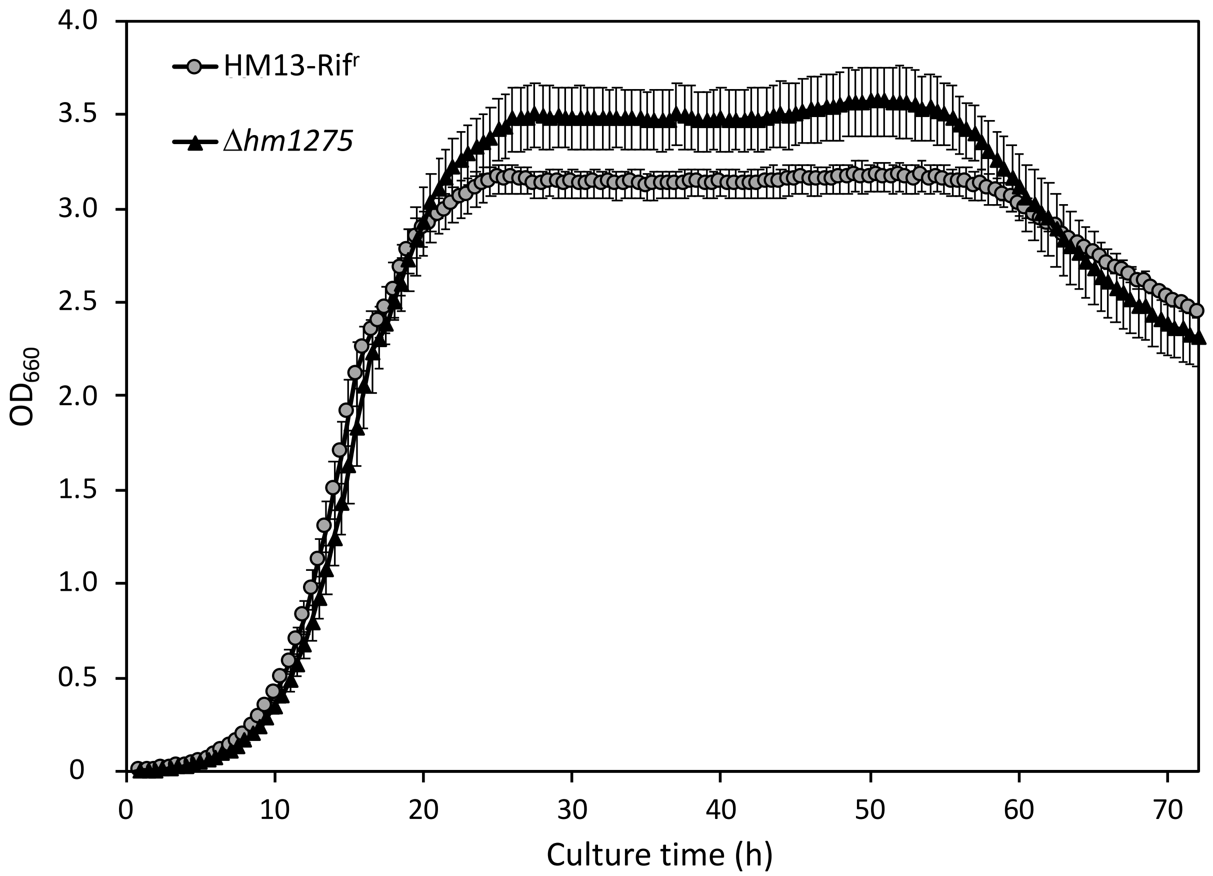


**Figure S2.** Growth of HM13-Rif$^r$ and Δ*hm1275* cultured for up to 72 h in M79 medium supplemented with CA. The data are the means ± standard errors of the values from three independent batches. Statistical analysis was performed using two-tailed unpaired Student's *t*-test, and no significant differences in growth were observed between the two strains at all time points ($p \geq 0.1$).

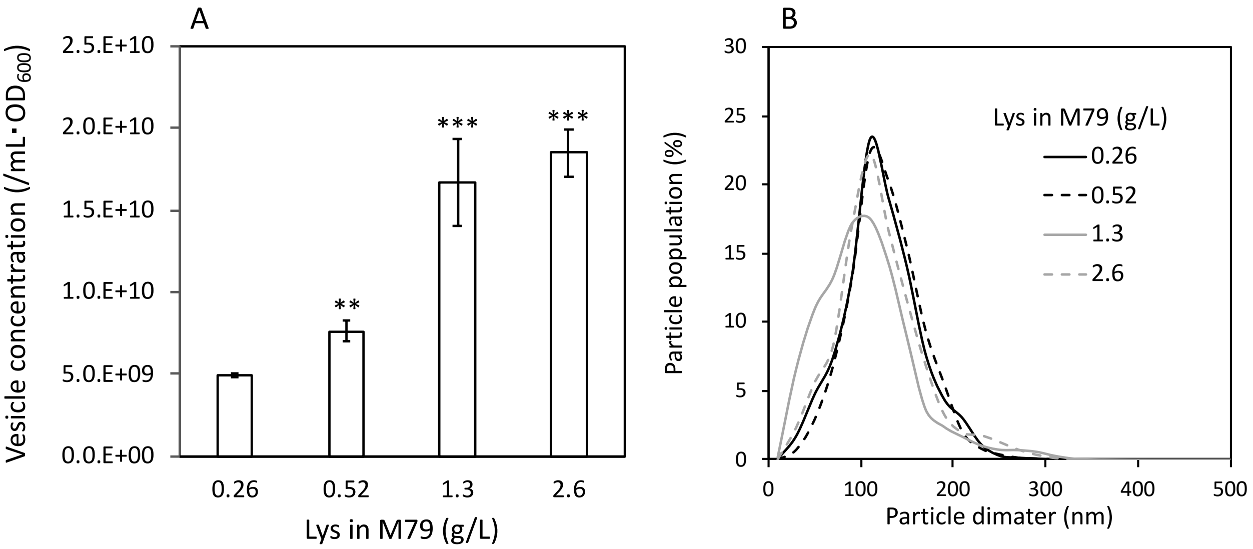


**Figure S3.** Evaluation of vesicle production of *S. vesiculosa* HM13 induced by the addition of Lys in a concentration-dependent manner by nanoparticle tracking analysis. Vesicle concentration (A) and particle size distribution (B) of HM13-Rif$^r$ cultured at different Lys concentrations. (A) The data are the means ± standard errors of the values from three independent batches. Statistical analysis was performed using two-tailed unpaired Student's *t*-test. ** and *** indicate $p < 0.05$ and 0.01, respectively, compared with the characteristics of vesicles produced in M79 medium supplemented with 0.26 g/L Lys. (B) All curves are the average from three independent batches.

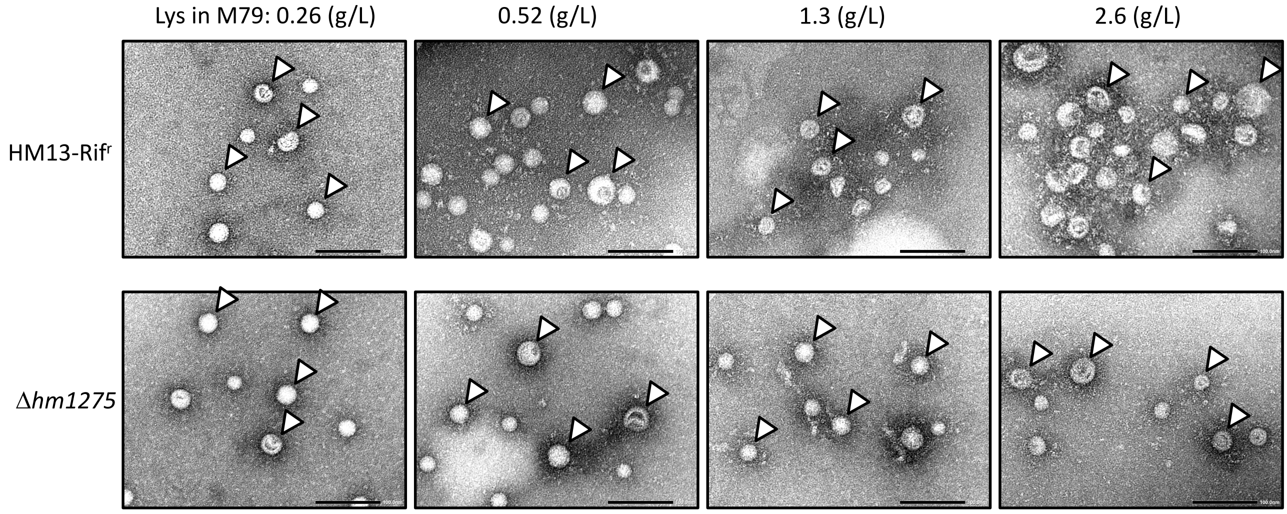


**Figure S4.** Similar morphology of EMVs produced by HM13-Rif[r] and Δ*hm1275*. Transmission electron microscopic micrographs of EMV fractions of HM13-Rif[r] and Δ*hm1275* cultured in modified M79 media with different Lys concentrations. The white arrowheads indicate representative EMVs. The bars indicate 100 nm.

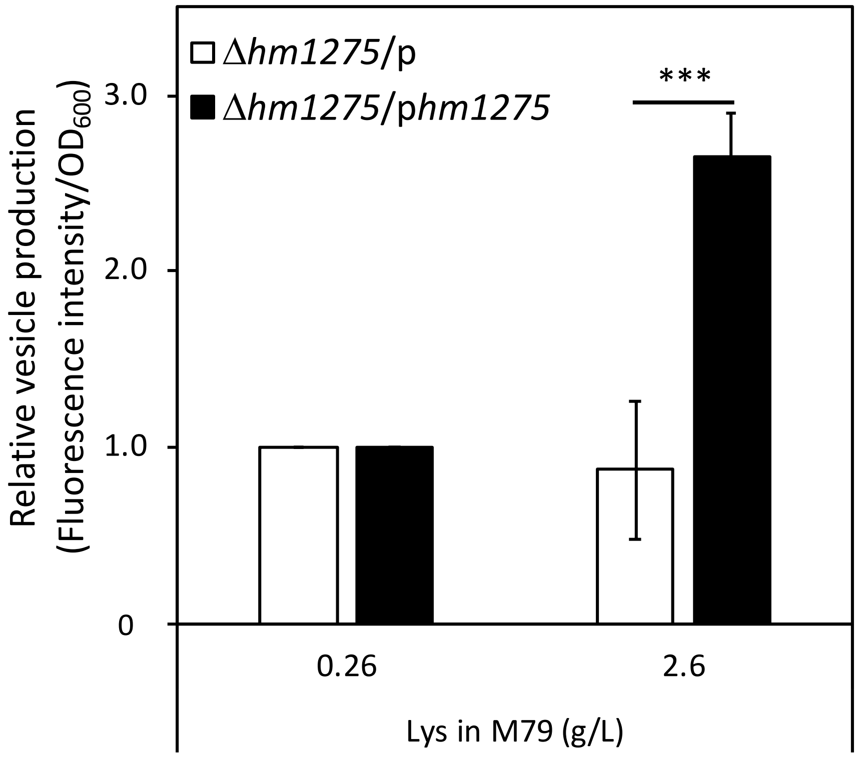


**Figure S5.** Complementation assay to demonstrate the role of HM1275 in Lys-induced vesicle production. The vesicle production of Δ*hm1275*/p and Δ*hm1275*/p*hm1275* was quantified by lipid staining. Each value of vesicle production was divided by the absorbance of cells grown in M79 containing 0.26 g/L Lys to compare relative vesicle production in each condition. The data are the means ± relative standard errors of the values from three independent batches. Statistical analysis was performed using two-tailed unpaired Student's *t*-test. *** indicates $p < 0.01$.

```
HM1275      MLMFNGRLKQENEQLKEELFSMQQVRKSLDSEMISMTLAADGKVTEVNQHFTNDMLFQQH     60
BdlA        -------------------------MAALDRSMARVEFDPDGNITDANENFLTLLGYRRD     35
                                       :** .*  : :  **::*:.*::* . : :::.  PAS1

HM1275      SPIGQQLIDMVPQRARGTAHYKSMCTAIKQSQHWAGALQIIKGNGEEAWLRAILQPVKKS     120
BdlA        EILGKPHRQLCDGAYAQSEDYRRFWERLRRGEHFSGRCKRITREGRPLWLEATYNPVRDG     95
            . :*:   ::       : .*: :   :::.:*::*  : *. :*.  **.*  :**:..  PAC1

HM1275      SGEVKYFSIYANDLTRTIETSKEHEDLIQALQRSTAVIEFDLNGTVLTANDNFLNAMGYK     180
BdlA        QGRLLKVVKYASDIDAIVHQEHEMQSKLDALSRSMAMIEFDLDGNVLAANDNFLATMGYG     155
            .*.:  .  **.*:   :. .:* :. ::**.** *:*****:*.**:****** :***   PAS2

HM1275      KNQLVGKHHKMFCEPQEYNSTEYQHFWEKLRNGNFVADRFKRVDSYGRVVWLEASYNPIF     240
BdlA        RAELASANHRQFCEPGYRDGPQYADLWRRLNRGEYVTGQFRRVHRNGQPVWLEASYNPVY     215
            : :*.. :*: ****   :. :* .:*.:*..*::*:.:*:**.  *: *********::  PAC2

HM1275      DSYNKLYKVVKFATVITDQVNREQAISEAADIAYSTSQQTDITAQRGNKVLLEMVEVMNK     300
BdlA        DADGKLYKVVKFASDVSDRMRRYQAEADNAHQAHTLSTETRTVAEHGALIIQSAVEEMLK     275
            *: .*********: ::*::.* ** :: *. *:: * :*  .*::*  :: . ** * *  MAT

HM1275      LALQMNAAAQGIEDLDKQSQMIGALVQSISGIADQTNLLALNAAIEAARAGEQGRGFAVV     360
BdlA        IANTLDASSLNIGELSQHSQQITSIVNTIREIAEQTNLLALNAAIEAARAGDQGRGFAVV     335
            :*  ::*:: .* :*.::** * ::*::*  **:*****************:********

HM1275      ADEVRQLASRTSKATEEIVEVVSKNKKLTENAVSMIEAGKKQALQGLTLSTQSGEVMSEI     420
BdlA        ADEVRQLAERTSKSTKEIADMIGRIQTGTRSVIDDMQHSQEQARRGVELANEAGAAILGI     395
            ********.****:*:**.:::.: :. *...:. :: .::** :*: *:.::* .:  *

HM1275      QDGAQKVVNAVGQFANQLNT--     440
BdlA        RESTHKVVEAVQQFSRTLNADL     417
            ::.::***:** **:. **:
```

**Figure S6.** Pairwise alignment of amino acid sequences of HM1275 and BdlA using Clustal Omega. These proteins shared a 38.8% identity. Colored underlining indicates the functional domains of BdlA shown in PDB. PAS: Per-Arnt-Sim, PAC: C-terminal region of several PAS sequences, MAT: methyl-accepting transducer. Asterisks (*) indicate identical residues, colons (:) indicate residues with strongly similar properties (STA, NEQK, NHQK, NDEQ, QHRK, MILV, MILF, HY, and FYW), and periods (.) indicate residues with weakly similar properties (CSA, ATV, SAG, STNK, STPA, SGND, SNDEQK, NDEQHK, NEQHRK, FVLIM, and HFY).

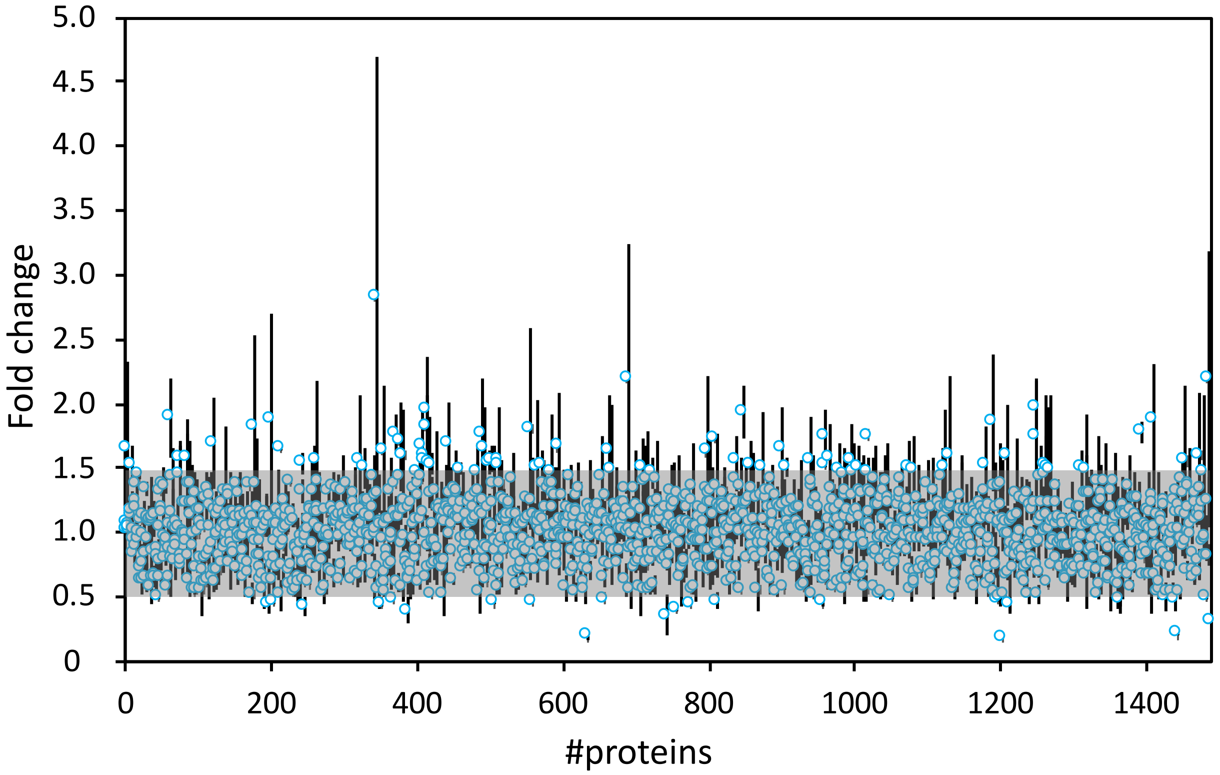


**Figure S7.** Proteins in *S. vesiculosa* HM13 cells up-/downregulated in response to Lys concentration in the medium. Proteins identified by shotgun proteomics were aligned according to the order of their gene location in the genome. The fold-changes were calculated by dividing the average expression values for 2.6 g/L Lys-grown cells by those of 0.26 g/L Lys-grown cells. The gray box indicates proteins with a fold change of 0.5–1.5. The data are the means ± standard errors of the values from two independent batches.

#### 3.2 Supplementary Tables

**TABLE S1** Estimation of functional domains in HM1275

| Name | Description | Range[a] | *E*-value[b] | Accession |
|---|---|---|---|---|
| **Signaling** | | | | |
| MCP signal | MCP (signaling domain) | 264–436 | $1.0\times 10^{-48}$ | cd11386 |
| MA | MCP-like domains (chemotaxis sensory transducer) | 252–436 | $1.8\times 10^{-45}$ | smart00283 |
| MCP signal | MCP (signaling domain) | 277–435 | $6.3\times 10^{-37}$ | pfam00015 |
| Tar | MCP | 248–440 | $2.0\times 10^{-36}$ | COG0840 |
| PRK15041 | MCP | 257–436 | $7.2\times 10^{-28}$ | PRK15041 |
| **Sensing** | | | | |
| PAS_3 | PAS fold | 166–253 | $2.6\times 10^{-12}$ | pfam08447 |
| PAS | PAS domain | 37–263 | $1.0\times 10^{-8}$ | COG2202 |
| PAS | PAS domain | 154–256 | $1.1\times 10^{-7}$ | cd00130 |
| Sensory box | PAS domain S-box | 156–266 | $6.1\times 10^{-7}$ | TIGR00229 |
| PAS_9 | PAS domain | 41–135 | $2.0\times 10^{-5}$ | pfam13426 |

| | | | | |
|---|---|---|---|---|
| PRK13560 | Hypothetical protein | 105–268 | $3.5 \times 10^{-3}$ | PRK13560 |
| PAS | PAS domain | 154–213 | $9.1 \times 10^{-3}$ | smart00091 |

[a] Range refers to the region of HM1275 (indicated by amino acid residue numbers) that shows sequence similarity to the domains provided.

[b] The *E*-value is the number of hits expected with a similar score by chance when searching the database with the amino acid sequence of HM1275.
